# Artificial selection for adult predation survival impacts life history and morphology in guppies (*Poecilia reticulata*)

**DOI:** 10.1101/2024.08.27.609842

**Authors:** Hannah De Waele, Regina Vega-Trejo, Catarina Vila-Pouca, Jori Noordenbos, Elizabeth Phillips, Bart J.A. Pollux, Alexander Kotrschal

**Affiliations:** Behavioural Ecology Group, Wageningen University & Research, 6708 WD Wageningen, The Netherlands; Department of Biology, Edward Grey Institute, University of Oxford, Oxford OX1 3SZ, UK; CEFE, Univ Montpellier, CNRS, EPHE, IRD, Montpellier, France

**Keywords:** life history, survival, guppy, reproduction, natural selection

## Abstract

Predation is accepted as a major evolutionary driver of life history and morphology. However, whether these traits evolve directly via predation or via indirect effects is largely unresolved. We used artificial selection over three generations to experimentally test the impact of adult predation on the evolution of life history and morphology in guppies (*Poecilia reticulata*). We found that, compared to control fish, predation-selected fish produced larger offspring and larger broods early in life. However, other life history parameters, such as interbrood interval and total number of offspring, showed no response. We also found that predation-selected for smaller and lighter females and for shorter tails and gonopodia in males, with no effect on body colouration. Our results show that while several traits evolve fast under selection on adult predation, several ‘classic’ predation-dependent traits seem unaffected by predation selection. By comparing our experimental results to those from natural populations we can disentangle the contribution of direct and indirect effects on trait evolution under predation pressure.

## Introduction

Predation is a key driver of evolutionary adaptations, as most animals face the risk of predation at some stage of their life. This threat from predators forces prey to balance seeking sustenance and mates, with avoiding becoming food themselves. As a result, prey species can undergo numerous changes to counteract predation risk. In evolutionary terms, animals can adapt to predation pressure by modification of their morphology or physiological performance (Caro, 2005), and/or adjustment of reproductive life history parameters (Stearns, 1992). For instance, adaptations in colouration and body shape can enhance a species’ ability to avoid detection, increase escape performance, or reduce their vulnerability to predators (Endler, 1980; Martin et al., 2014; Young et al., 2011). Examples include chameleons evolving camouflage to blend in with their surroundings (i.e. *Bradypodion taeniabronchum*) (Stuart-Fox et al., 2008), frogs evolving bright colouration to signal toxicity (i.e. *Dendrobatidae*) (Davies et al., 2012), or fish evolving spines as defensive structures (i.e. *Gasterosteus aculeatus*) (Reimchen 1992, Barret 2008). Some species may also evolve heightened sensory capabilities to facilitate the detection of potential predators; for example, the crustacean *Gammarus minus* evolves larger eyes in high-predation environments (Glazier & Deptola, 2011). Animals can also adopt changes in body size and shape to enhance escape performance in the presence of predators (Hendry et al., 2006; Langerhans et al., 2004; Walker et al., 2005). Western mosquitofish (*Gambusia affinis*), for example, exhibit body shape differences that allow for faster burst speed to facilitate escaping predators in high-predation environments (Langerhans et al., 2004).

In addition to such common morphological adaptations aimed at increasing survival in the face of predation (Caro, 2005), predation can have a profound impact on the evolution of life history strategies of prey species (Arendt & Reznick, 2005; Martin, 1995; Reznick, 1982; Reznick et al., 1990, 1996, 2001; Reznick & Bryga, 1987, 1996; Reznick & Endler, 1982; Riesch et al., 2013; Stearns, 1989). An animal’s life history strategy is defined by the collection of variables that contribute to an individuals’ lifetime reproductive success (Roff, 1993; Stearns, 1989), such as age at first reproduction, investments in reproduction versus somatic maintenance, and offspring characteristics (small vs large; few vs many) (Losos, 2017). Life history theory predicts evolution to drive an optimal allocation of resources that results in the largest number of successful offspring best suited to the environmental conditions (Roff, 1993; Stearns, 1989, 1992) and predation modulates these strategies. For instance, prey often shift their growth rate and age of maturity when predation risk is elevated; the freshwater snail *Physella virgaga virgata* exhibits higher age and size at maturity in the presence of a predator crayfish (*Orconectes virilis*) (Crowl, 1990; Crowl & Covich, 1990) as crayfish prey selectively on the smallest snails. This causes high mortality in juveniles, while adults appear to have refuge due to their size. In some mammal (Creel et al., 2007; Korpimaki et al., 1994) and bird species (Dillon & Conway, 2018; Eggers et al., 2006; Zanette et al., 2011) elevated predation risk concurs with fewer offspring per reproductive event, while certain fish and invertebrate species often produce more and smaller offspring under increased predation pressure (Johnson & Belk, 2001; Reznick, 1982; Stibor, 1992). This is due to species-specific ecological constraints, which underlie species-specific responses in life history evolution, that allow species to fine-tune their reproductive strategy to maximize fitness under high-predation risk.

Much of our current knowledge on predation-driven adaptations in life history and morphology stems from research on the Trinidadian guppy (*Poecilia reticulata*) (Bronikowski et al., 2002; Endler, 1980, 1991; Houde, 2019; Magurran, 2005; Reznick, 1982; Reznick et al., 2001; Reznick & Bryant, 2007). This is because guppies inhabit parallel streams that naturally exhibit a range of predation pressures: downstream areas are typically home to several predator species (e.g. *Crenicichla frenata*, *Hoplias malabaricus*) that prey on adult guppies (high-predation environments), while waterfalls block access of most predators to upstream regions that therefore have few to no such predators (low-predation environments). These replicated predation differences allow us to compare natural guppy populations that evolved under different levels of predation pressure. Indeed, guppies from high and low-predation environments exhibit a range of differences in many morphological and life history traits (Houde, 2019; Magurran, 2005). For example, male guppies in low-predation habitats develop more vibrant colouration compared to fish from high-predation habitats (Endler, 1978). This phenomenon is attributed to the decreased selection pressure for crypsis due to fewer predators, while sexual selection, driven by female preference for conspicuous males, remains strong. In line with this, relocation of high-predation guppies to low-predation environments leads to rapid evolution of colourful males (Endler, 1978, 1980a; Gordon et al., 2015; Gotanda et al., 2019; Kemp et al., 2018). Predation-dependent effects on body shape have also been reported in guppies (Alexander et al., 2006; Burns et al., 2009; Hendry et al., 2006; Langerhans & DeWitt, 2004), with natural selection likely favoring an improved fast-start escape performance in high-predation environments since this is crucial in avoiding capture during a predator strike (Eaton, Bombardieri, and Meyer 1977; Harper and Blake 1990; Howland 1974; Webb 1978; Weihs 1973). Concurrently, in the closely related genus *Gambusia* (family Poeciliidae), predation risk seems to drive the evolution of morphological features that facilitate an increased fast-start escape velocity, such as an elongated body and smaller head size to increase streamlining (Langerhans et al. 2004), and a shorter gonopodium (modified anal fin) to reduce drag (Langerhans, Layman, and DeWitt 2005). However, in guppies, evidence is not congruent on how body shape and fin size evolve in high-predation contexts (Burns, Di Nardo, and Rodd 2009). In addition to the effects of predation on morphological evolution, the guppy system has been key in developing and testing concepts of life history evolution (Stearns 1989). It is generally assumed that natural selection should favour maternal investment to produce larger numbers of offspring at a relatively rapid pace in high mortality environments, compared to environments with low mortality rates (Charlesworth 1994; Gadgil and Bossert 1970; Law 1979; Michod 1979). Indeed, previous research shows that, compared to low-predation guppies, guppies from high-predation habitats mature at an earlier age and at a smaller size, produce larger broods with smaller offspring, have shorter interbrood intervals (defined as the number of days between two births), and invest more into reproduction early in life (Reznick 1982, 1989; Reznick and Bryga 1996; Reznick and Endler 1982; Reznick, Rodd, and Cardenas 1996).

Studying the effect of predation on trait evolution through the comparison of natural populations introduces the complexity of varying ecological characteristics beyond differences in predation pressure. For instance, when comparing wild populations of guppies, streams with high-predation pressure also tend to be characterized by lower species richness, lower conspecific density, wider channels, more open canopies, and higher levels of primary productivity (Feminella, Power, and Resh 1989; Hawkins, Murphy, and Anderson 1982; Magurran 2005; Power 1984; Reznick and Endler 1982). Hence, when comparing traits of fish from low and high-predation populations the *direct* effect of predation is inherently confounded by *indirect* effects of confounding ecological traits, that are likely to also influence the measured traits. For instance, higher primary production in high-predation pressure populations may lead to shorter interbrood intervals simply because of higher food availability. While studies comparing natural populations have been instrumental in uncovering broad evolutionary patterns and generating hypotheses, they do not allow for causal conclusions about the effects of predation on trait evolution. Therefore, we here introduce an experimental approach to establish causality between predation pressure and trait evolution. In this study, we experimentally test the effect of adult predation on the evolution of life history and morphological traits in guppies. We set up an artificial selection experiment based on adult survival during brief periods of cohabitation with a predator (*Crenicichla alta*) over three consecutive generations. To do this, we established three predation survival lines and three control lines. We focused on the effect of direct removal by predators while controlling for potential foraging effects by providing high food availability. Additionally, we controlled for non-lethal effects of visual and olfactory predation cues by keeping the control fish in the same tank, but protecting them from being consumed with a translucent, water-permeable barrier.

Based on previous correlative evidence on predator-driven morphology evolution (Alexander et al. 2006; Burns, Di Nardo, and Rodd 2009; Endler 1980; Gotanda et al. 2019; Hendry et al. 2006; Kemp, Batistic, and Reznick 2018; Langerhans and DeWitt 2004), we predicted that predation would select for smaller adults, shorter gonopodia, larger eyes, and less colourful males and possibly a combination of body shape or caudal fin size that allows for a fast escape. Additionally, based on life history evolution theory (Stearns 1992) and correlative evidence (Reznick 1982; Reznick and Endler 1982; Reznick, Rodd, and Cardenas 1996), we predicted that predation would select for higher fecundity, more frequent reproduction or shorter interbrood interval, and smaller offspring size. Any deviations from those commonly accepted differences would then indicate that factors besides adult predation were underlying previously observed trait evolution and therefore merit reconsideration of such supposedly established relationships.

## Results

### Artificial selection in adult guppies

We examined the effect of adult predation on reproductive and morphological traits by comparing fish from populations that were subjected to divergent experimental predation pressures. In brief, we placed 212 fish into a large semi-natural tank designed to mimic the natural habitat of guppies including a pike cichlid as predator. Thirty-two randomly selected fish were placed behind a barrier (safe from consumption), while the remaining 180 were monitored daily until approximately 80 % were consumed (after 45.6 days ± 28.3 in the tank 81% ± 3 animals consumed). In the three ‘control’ groups all animals survived the phase of cohabiting with the predator behind a barrier. Males and females were run separately and paired up to produce the next generation. Once their progeny matured, they was subjected to the same selection procedure. We did this for three generations and three replicates and then assessed the reproductive traits of the third-generation fish (F3; Figure 1), and the morphological traits of their offspring (F4). Details on the artificial selection procedure can be found in the Materials and Methods section.

**Figure 1.**
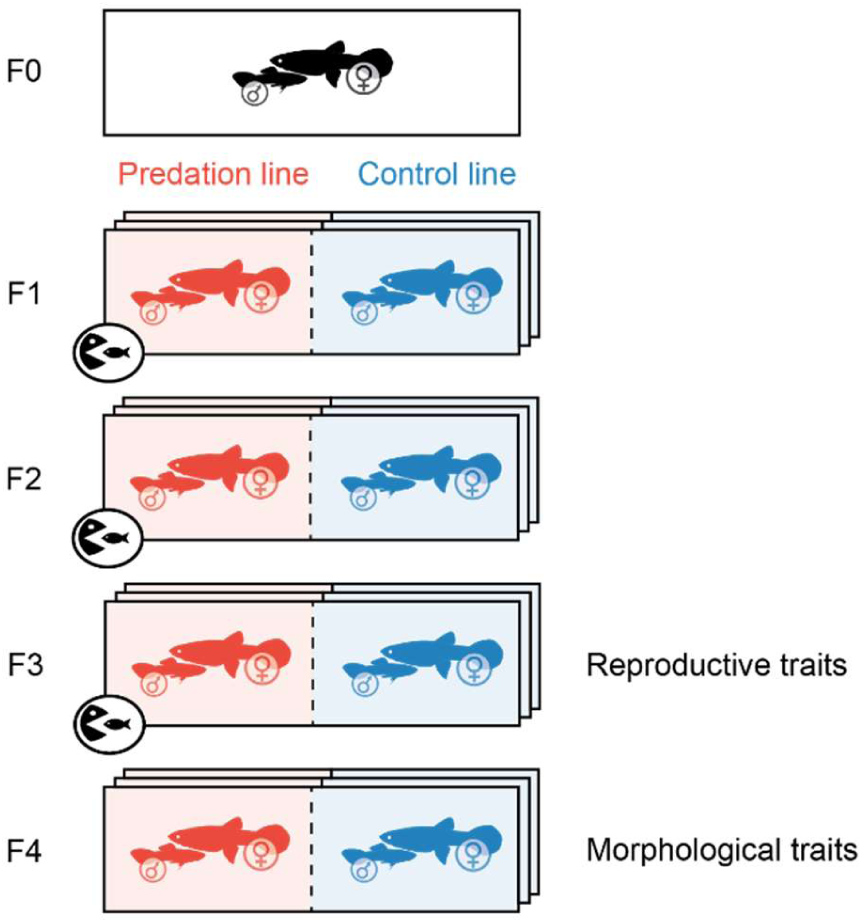
Overview of the generations of fish used throughout this experiment. Fish from generation F1, F2, and F3 were subjected to a predator (predation fish – in red – underwent selection for survival under predation, while control fish – in blue – were subjected to sensory cues only). Fish from generation F3 were examined for differences in reproductive traits, while fish from generation F4 were tested for differences in morphological traits. Each generation had three replicates.

### Predation impacts various reproductive traits

We found that fish from the predation and control treatments (F3) did not differ in their likelihood to breed (Table 1; est = 18.6, se = 4644, Z = 0.004, p = 0.997), produced their first brood at comparable times from first coupling (Table 1, est = −0.02, se = 0.03, Z = −0.71, p = 0.478), and produced a similar total number of offspring (Table 1, est = −0.006, se = 0.07, Z = −0.076, p = 0. 940). However, when producing their first broods, predation females produced larger broods than control females, but with consecutive broods they went on to produce smaller broods. (Table 1; treatment: est = 0.19, se = 0.08, Z = 2.46, p = 0.014; brood: est = - 0.06, se = 0.02, Z = −4.34, p < 0.001; treatment x brood: est = −0.07, se = 0.02, Z = −3.25, p < 0.001; Figure 2).

**Figure 2.**
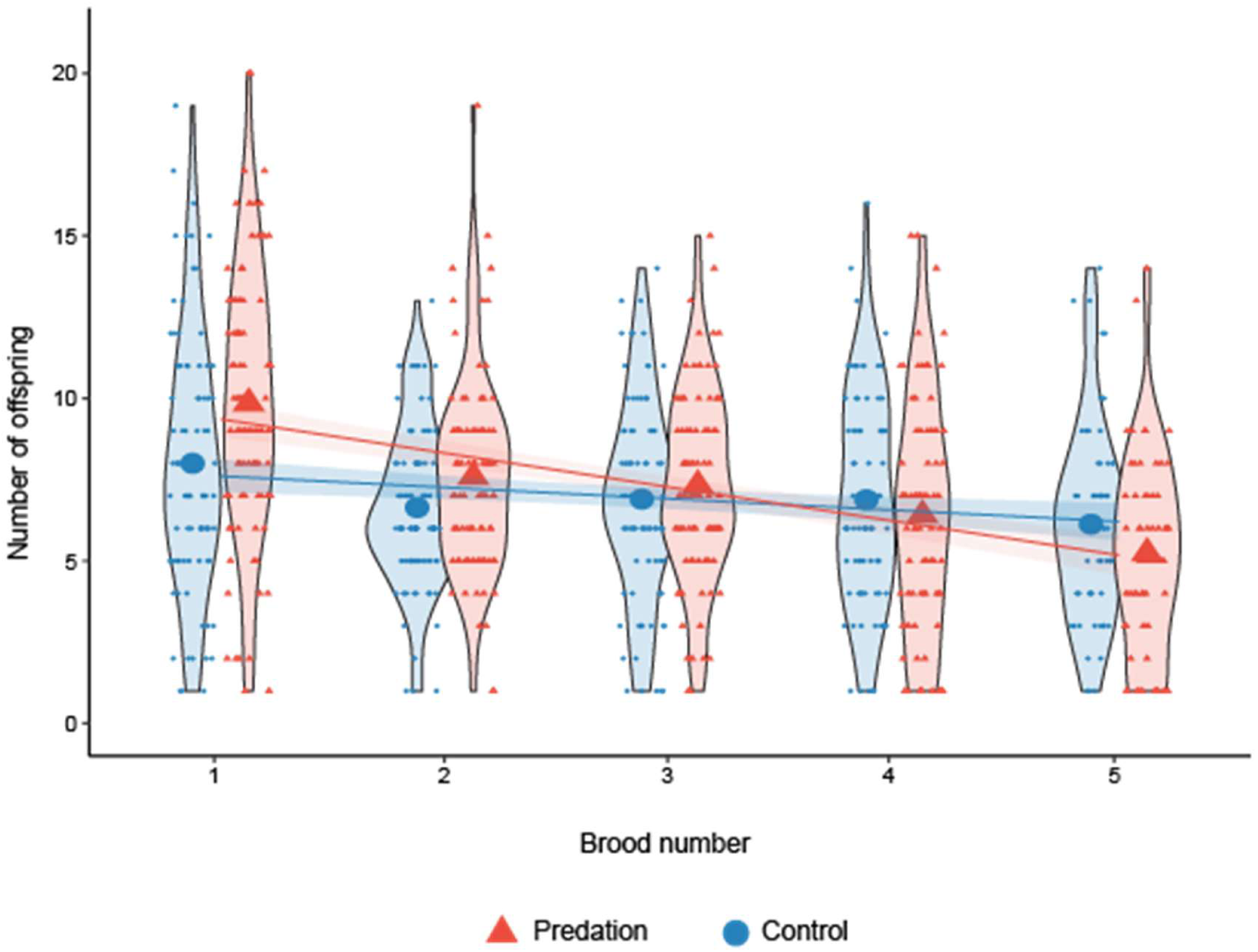
Predation females produced larger first broods compared to controls. Differences in number of offspring over subsequent broods between predation (red triangles) and control (blue circles). Model predictions are plotted as the best fit line and shaded areas indicate the SE. Violin plots show the distribution of the data.

**Table 1.** Descriptive characteristics of the study groups (Predation and Control) with means ± SD, and p-value for each of the characteristics. Dark red background indicates the highest value, while light red background indicates the lowest value.

| Characteristic | Predation |  | Control | P-value |
| --- | --- | --- | --- | --- |
| **F3** |  |  |  |  |
| Offspring production | 96 (100%) | = | 93 (97.9%) | 0.997 |
| *Number of offspring per brood* | 7.46 ± 3.64 | > | 6.89 ± 3.18* | **0.014** |
| Total number of offspring across all broods | 33.8 ± 9.85 | = | 32.1 ± 11.5 | 0.940 |
| Total amount of broods | 4.53 ± 1.00 | = | 4.66 ± 1.04 | 0.984 |
| Time to first brood | 53.7 ± 13.3 | = | 55.1 ± 13.0 | 0.478 |
| Interbrood interval | 41.7 ± 13.2 | = | 41.5 ± 12.9 | 0.275 |
| *Standard length (mm) of juveniles* | 7.97 ± 0.45 | > | 7.62 ± 0.35* | **0.005** |
| **F4** |  |  |  |  |
| Standard length (mm) |  |  |  |  |
| **Females** | 26.4 ± 2.13 | < | 27.9 ± 2.21* | **< 0.001** |
| Males | 20.6 ± 1.35 | = | 20.6 ± 1.19 | 0.752 |
| Relative weight (mg) |  |  |  |  |
| **Females** | 421 ± 117 | < | 446 ± 108* | **< 0.001** |
| Males | 157 ± 30.6 | < | 164 ± 29.9 | 0.153 |
| Relative body area - bulkiness (mm<sup>2</sup>) |  |  |  |  |
| **Females** | 140 ± 25.1 | < | 147 ± 31.7* | **0.005** |
| Males | 79.8 ± 10.8 | = | 80.7 ± 10.7 | 0.306 |
| Relative tail length (mm) |  |  |  |  |
| Females | 8.17 ± 0.60 | = | 8.41 ± 0.75 | 0.441 |
| **Males** | 6.97 ± 0.77 | < | 7.33 ± 1.04* | **< 0.001** |
| Relative eye size (mm<sup>2</sup>) |  |  |  |  |
| Females | 3.53 ± 0.43 | = | 3.73 ± 0.39 | 0.383 |
| Males | 2.33 ± 0.30 | = | 2.29 ± 0.27 | 0.065 |
| *Relative gonopodium size (mm)* | 3.75 ± 0.42 | < | 4.24 ± 0.45* | **< 0.001** |
| Relative male colouration (mm<sup>2</sup>) |  |  |  |  |
| Black | 13.1 ± 4.38 | = | 13.3 ± 4.22 | 0.743 |
| Orange | 4.73 ± 3.06 | = | 4.43 ± 3.01 | 0.346 |
| Iridescence | 5.05 ± 4.73 | = | 5.20 ± 4.22 | 0.795 |

Over the course of the 37 weeks we recorded breeding metrics, neither the number of broods produced (Table 1; est = 0.002, se = 0.10, Z = 0.02, p = 0.984), nor the interbrood intervals differed between treatments (Table 1; and est = −0.05, se = 0.05, Z = −1.09, p = 0.275). Yet, newborn offspring from the predation treatment were larger than control newborns (Table 1 and Figure 3; est = 0.40, se = 0.14, Z = 2.80, p = 0.005).

**Figure 3.**
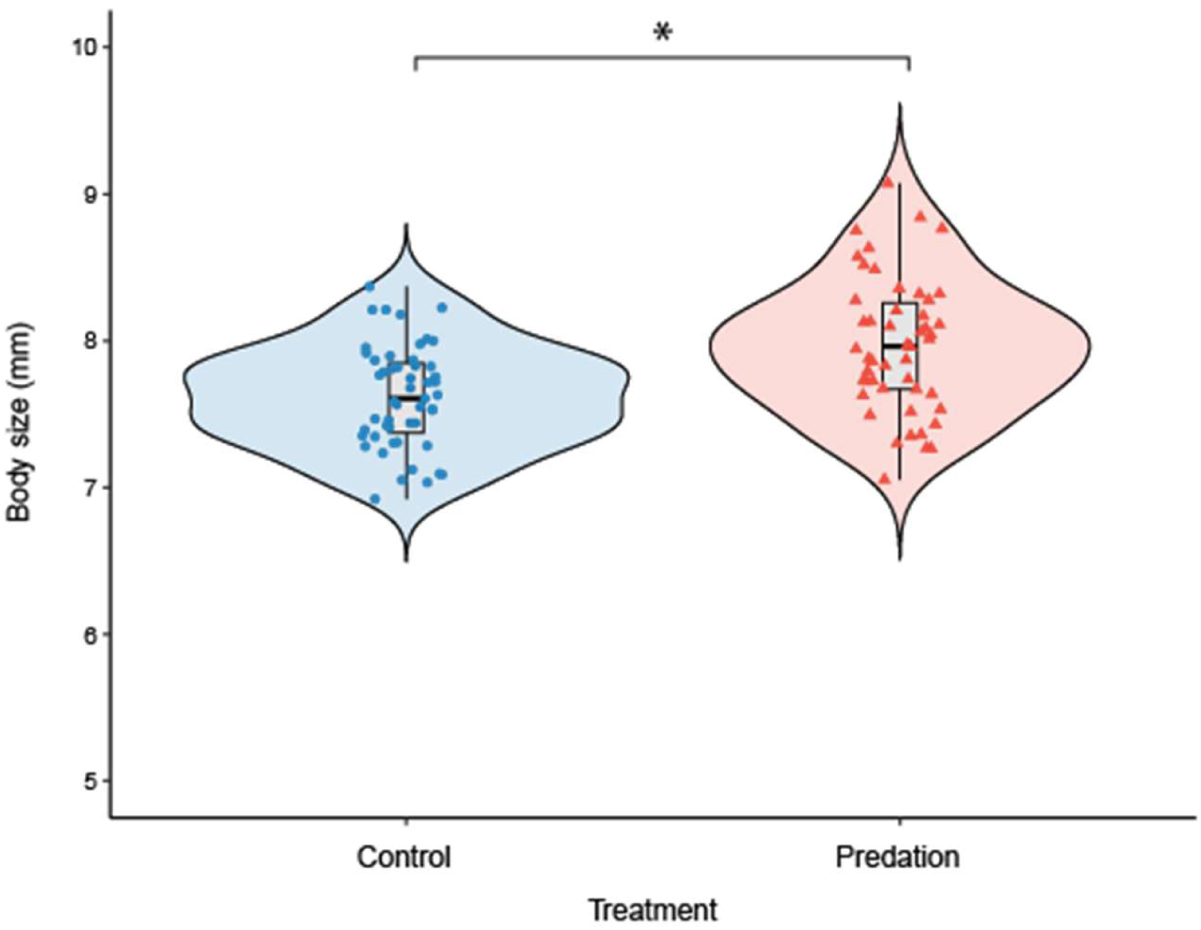
Offspring from the predation treatment were larger than controls. Raw data for body size for juveniles for control (blue circles) and predation (red triangles) treatment for F4 generation. The grey box indicates the median and interquartile range (IQR), whiskers extend 1.5*IQR. Violin plots show the distribution of the data.

### Predation impacts several morphological traits

We found sex-dependent effects of predation on animals whose parents, grandparents and great-grandparents were subject to intense adult predation selection. Female offspring of predation group animals were smaller (Table 1 and Figure 4a; est = −1.43, se = 0.25, Z = −5.74, p < 0.001) and lighter (Table 1 and Figure 4a; est = −396, se = 114, Z = −3.50, p < 0.001) than control females. We also found a steeper allometry in predation females’ body mass: smaller females were lighter, and larger females were heavier compared to similar sized females from control lines (Table 1 and Figure 4a; treatment x standard length: est = 15.56, se = 4.16, Z = 3.74, p < 0.001). Additionally, predation females were more ‘streamlined’ as their relative body area was smaller (Table 1; est = 5.81, se = 2.09, Z = 2.78, p = 0.005) compared to control females. However, we found no differences in tail length (Table 1; est = 0.05, se = 0.06, Z = 0.77, p = 0.441), nor eye size (Table 1; est = −0.04, se = 0.04, Z = −0.87, p = 0.383) for females between treatments.

**Figure 4.**
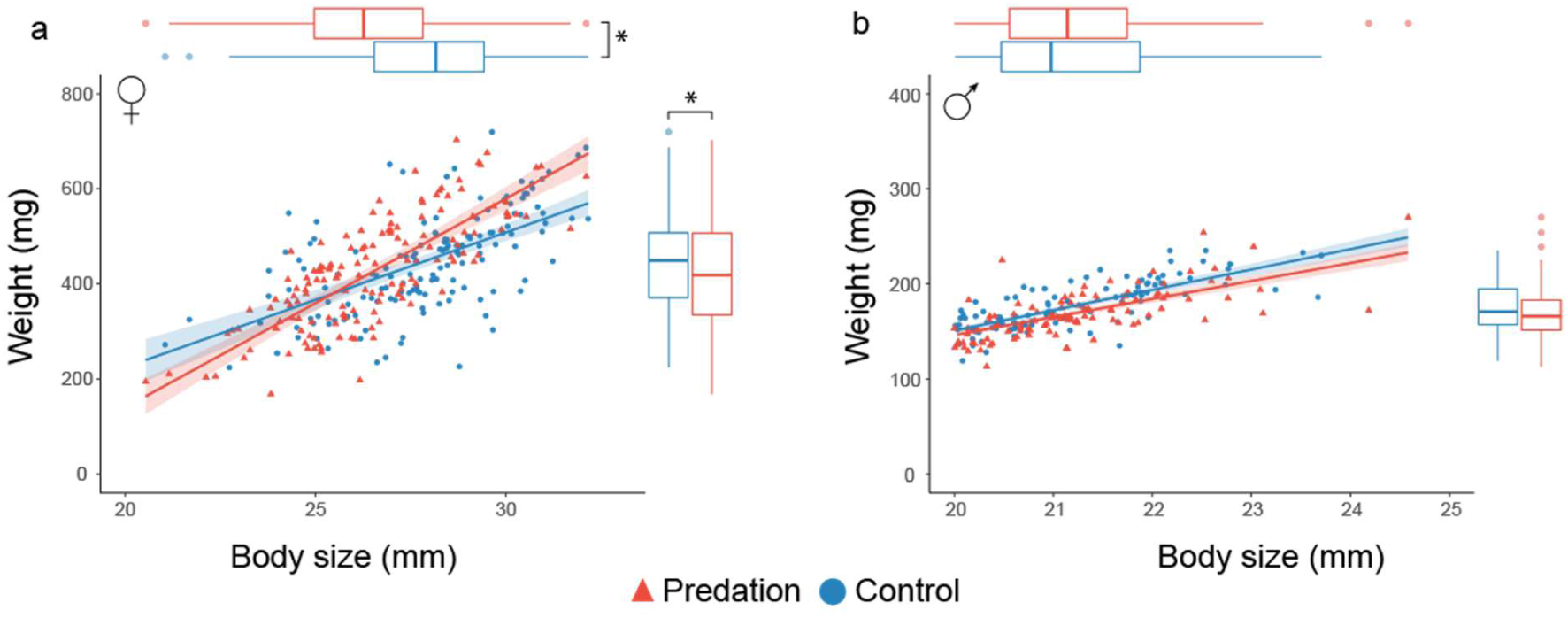
Offspring of the predation treatment has a steeper body mass allometry compared to controls. Raw data for weight compared to body size in F4 generation (a) females and (b) males between control (blue circles) and predation (red triangles) for females. Model predictions are plotted as the best fit line and shaded areas indicate the SE. Boxplots for both weight and body size indicate the median and interquartile range (IQR), whiskers extend 1.5*IQR, and data beyond the end of the whiskers are plotted individually.

For males, we found no differences between treatments in either body size (Table 1 and Figure 4b; est = −0.05, se = 0.14, Z = −0.32, p = 0.752), body weight (Table 1 and Figure 4b; est = 45.0, se = 31.5, Z = 1.43, p = 0.153), or streamlining (i.e. relative body area (est = −0.51, se = 0.50, Z = −1.02, p = 0.306)). However, predation males evolved shorter tails (Table 1; est = - 0.36, se = 0.10, Z = −3.58, p < 0.001) and shorter gonopodia (Table 1; est = −0.49, se = 0.05, Z = −9.72, p < 0.001). We found no pronounced differences in eye size between treatments (Table 1; est = 0.05, se = 0.03, Z = 1.84, p = 0.065). Lastly, we found no differences in colouration between predation and control groups in relative area of black, orange, or iridescence (Table 1; black: est = −0.15, se = 0.45, Z = −0.33, p = 0.743; orange: est = 0.32, se = 0.34, Z = 0.94, p = 0.346; iridescence: est = −0.13, se = 0.49, Z = −0.26, p = 0.795). In each model, the replicates are controlled for; full output of all models can be found in the supplemental information.

## Discussion

Our study shows that after three generations of adult predation, predation-selected and control guppies differ in some of the morphological and life history traits commonly associated with predation evolution. It is unlikely that these effects are driven by confounding factors often covarying with predation pressure as our experiment was set up to minimize those. Our results therefore confirm, to our knowledge, for the first time experimentally, that predation survival is indeed causally linked to the evolution of offspring size and number, adult body size and shape, and gonopodium size.

Theory predicts that increased adult mortality due to predation causes evolutionary shifts in life history traits in prey (Charlesworth 1994; Gadgil and Bossert 1970; Law 1979; Michod 1979). Natural selection should favor mothers to produce larger numbers of offspring at a relatively rapid pace in high mortality environments, compared to environments with low mortality rates. This is supported by prior empirical evidence showing that animals from high-predation environments produce more, but smaller offspring with shorter interbrood intervals and that they invest a higher proportion of consumed resources to reproduction early in life (Evans, Gasparini, and Pilastro 2007; Reznick 1982, 1989; Reznick and Endler 1982). Our findings partially confirm this theory, showing that prey under high-predation pressure invest in producing more offspring early in life. However, we did not observe an increase in the frequency of reproduction or in the total number of offspring. Our results suggest a shift in reproductive investment towards early offspring production and are in line with the prediction that predation selects for increased early reproductive investment (Williams 2001).

Conversely, we did not find the traditional trade-off between offspring number and size (D. A. Roff, Mostowy, and Fairbairn 2002; Smith and Fretwell 1974; Stearns 1992). We even found an opposite effect in investment of juvenile size where predation lines had more and larger offspring. While field data consistently reports newborns that are about half as heavy in high-predation compared to low-predation localities (Dial, Reznick, and Brainerd 2016; Reznick 1982; Reznick and Endler 1982; Reznick, Rodd, and Cardenas 1996), we found a small increase in newborn size in predation compared to control line newborns. This suggests that the patterns found in previous empirical studies are a result of other agents of selection acting in concert with predation pressure. For example, low-predation localities are known to have higher densities of conspecifics (Reznick et al. 2012) and females produce larger offspring at birth in response to these increased densities (Leips et al. 2009; Leips, Helen Rodd, and Travis 2013). Additionally, density affects food availability, with low-predation localities having lower food availability, which can also influence juvenile size. This was exemplified in an experiment where isolated adult female guppies were maintained on either low or high food rations and responded to low food rations by producing larger offspring (Reznick and Yang 1993). These mechanisms could explain the increased juvenile size found in studies in natural populations. Additionally, when considering predation pressure, we would expect predation to select for larger offspring, as bigger is generally better for their survival (Einum and Fleming 2000; Marshall and Keough 2008; Wilson et al. 2005). Larger offspring are more competitive, experience a shorter vulnerable juvenile period, and thereby enjoy higher survivorship (Bashey 2008; Falster, Moles, and Westoby 2008; Herrel and Gibb 2006; Wassersug and Sperry 1977; Werner and Gilliam 1984). The main threat to guppy survival in high-predation sites is imposed by larger fishes (O’Steen, Cullum, and Bennett 2002), and when attacked by a predator, guppies, like most fishes, attempt to escape by performing a rapid swimming behaviour: the fast-start escape response. Faster starts have been shown to increase the probability of adult guppies escaping from natural predators (Walker et al. 2005), thus resulting in higher survival (Dial, Reznick, and Brainerd 2016; O’Steen, Cullum, and Bennett 2002), and larger prey fish are expected to show higher escape performance due to larger muscle mass and body length (Wakeling, Shadwick, and Lauder 2006; Webb 1976). It is noteworthy that in our experiment, we focused on adult predation, while in a high-predation environment animals can be exposed to predation during all life stages. As antipredator strategies can differ between juveniles and adults, additional experiments also focusing on juvenile predation as a selective agent may be a fruitful avenue for future research.

In the adults, we found sex-dependent differences in body size and shape. In females, predation fish became smaller, less heavy, and more streamlined than the control fish. Body size differences could be explained by two non-mutually exclusive processes, a hereditary effect of size-selective predation if the pike cichlid preferred to prey on larger females (Johansson, Turesson, and Persson 2004) and/or strong selection for maturing early in high-predation environments. To disentangle those effects and conclusively clarify the mechanism by which females that survive predation are smaller than control fish, growth rate parameters and age at maturity data will be necessary. In addition to body size, body shape is often related to predation-dependent selection pressures. In general, predation environments select for a more streamlined body and deeper caudal peduncles, which presumably increases escape ability and survival (Gomes and Monteiro 2008; Langerhans et al. 2004; Langerhans and Makowicz 2009). For example, in several Poecilid fishes, body streamlining through superfetation has been shown to improve the fast-start escape performance (Fleuren, van Leeuwen, and Pollux 2019). In line, we found that predation fish have less body area when controlling for length. As this is only a proxy for streamlining, additional studies in geometric morphometrics complemented with swim tunnel assays will be necessary to clarify this matter. In males, however, we found support for streamlined phenotypes where fish from the predation treatment had shorter caudal (tail) and anal (gonopodium) fins than controls. While increased predation is at times correlated with shorter caudal peduncles (Langerhans and Makowicz 2009), to our knowledge data on caudal fin size vs. predation is currently lacking. Generally, selection for increased caudal fin area is thought to increase the speed of fast starts (Ghalambor, Walker, and Reznick 2003). Additionally, hydromechanical theory predicts that large fin area should be advantageous for fast-start performance (Weihs 1972, 1973). Indeed, a reduction of acceleration performance is measurable after amputation of (parts of) the caudal fin in Rainbow Trout (*Salmo Gairdneri*) (Webb 1977). Hence, a large caudal fin seems required for maximum acceleration performance. Parallel to this, a study in guppies shows that higher burst speeds increase the probability of survival of a predator attack (Walker et al. 2005). Additionally, guppies from high-predation localities are more likely to survive a predator attack (O’Steen, Cullum, and Bennett 2002; Seghers 1973). Yet, in our experiment, we find that high-predation environments favor *shorter* caudal fins. To elucidate this discrepancy, future assays of male swimming abilities will uncover whether smaller fins aid in escaping predation in guppies. Regarding the length of the gonopodium, evidence is unanimous that elaborate male traits favored by females often increase susceptibility to predation (Andersson 1994; Rosenthal et al. 2001). Additionally, long gonopodia increase drag and would inhibit burst speeds (Langerhans, Layman, and DeWitt 2005). Therefore, our findings that fish subject to predation have shorter gonopodia than control fish are in line with expectations. Yet, in studies of natural populations, high-predation environments often reveal larger gonopodia (Jennions and Kelly 2002; Kelly, Godin, and Abdallah 2000). This may be explained by the fact that males in these high-predation localities typically minimize active sexual display to females but resort to a higher proportion of sneak-copulations (Magurran and Seghers 1994a). Longer gonopodia are likely beneficial for such coercive mating tactics. Indeed, poecilids with sneak-copulations generally have longer gonopodia than species with male courtship (Jennions and Kelly 2002; Rosen and Tucker 1961). This can be explained by the fact that sneaking behaviour is less conspicuous to potential predators than courtship (Godin and Dugatkin 1995; Magurran and Nowak 1991; Magurran and Seghers 1990). Additionally, females in high-predation environments are more concerned with the threat of predation and are less vigilant towards sneaky mating attempts (Houde and Hankes 1997). In our experiment we removed any potential for sexual selection as we decided on mating partners and found the opposite relationship between gonopodium length and predation pressure. We attribute the shorter gonopodia of the predation-selected fish to an improvement in swimming ability - specifically burst speed.

Male colour phenotypes are subject to antagonistic sexual and natural selection; females prefer more conspicuous male phenotypes, but these also confer greater predation risk (Godin and McDonough 2003). Therefore, we expect the expression of this sexual trait to reflect a balance between natural and sexual selection (Andersson 1994; Houde and Hankes 1997; Reznick and Endler 1982). In line with previous predictions, we expected high-predation to select for less conspicuous colours and patterns (Endler, 1978, 1980a, 1984; Gordon et al., 2015; R. A. Martin et al., 2014; Millar et al., 2006). Yet, in our experiment, where predation was high and sexual selection was removed, we found no changes in colouration. While multiple studies, including classic ‘textbook’ examples (Endler 1980), have provided evidence for evolution in guppy colour patterns as a function of predation, several subsequent studies suggest that any such effects may be small. For example, guppies from high-predation populations transplanted in mesocosms without predators showed increased black spot sizes after a year but inconsistent changes in all other colour traits (Gotanda et al. 2019). Additionally, multiple guppy transplant studies have reported inconsistent differences across regimes and within rivers (Dick et al. 2018; Kemp et al. 2009; Kemp, Batistic, and Reznick 2018). Finally, when quantifying the perception of colour patterns as seen by both guppies and pike cichlids, a recent study concluded that predation had no consistent effect on the evolution of colouration in male guppies (Yong et al. 2022). Clearly, there are differences in colouration between low and high-predation populations in many drainages, but the question remains whether these phenotypic differences are driven by differences in predation. To accurately discern the role of predation on colour pattern evolution, differences among drainages must be controlled for, and Endler’s classic experiment did not explicitly focus on pairing high and low-predation populations and was, therefore, unable to do so (Endler 1978). May this indicate that predation is not the main driver for colouration in guppies, but female choice is. It is well established that females prefer conspicuous males (Houde and Hankes 1997). Yet, there is substantial geographic variation in female preferences for colour phenotypes (Endler and Houde 1995), which could drive large variation in male colouration. In addition, in high-predation environments, female preoccupation with predator evasion is higher (Magurran and Nowak 1991), allowing males to engage in higher rates of (successful) sneaky matings (Farr 1975; Luyten and Liley 1985; Magurran and Seghers 1994b; Matthews, Evans, and Magurran 1997). In such environments, sexual selection may become less important, eliminating the need for high conspicuousness to attract a mate. In our experimental set-up, we eliminated sexual selection completely by keeping sexes apart and assigning breeding partners, and we found no direct effect of predation on male colouration. We therefore conclude that increased predation on more colourful - supposedly for predators more salient - males is not a key driver of the evolution of guppy colouration.

Many selective agents could have left their imprint on trait evolution in studies with natural populations. In every study examining adaptations in natural populations, scientists conclude that adaptations represent responses and compromises to multiple agents of selection (Reznick, Butler IV, and Rodd 2001). Thus, when comparing wild guppies from different predation regimes to discern the role of predation on trait evolution, causally linking predation to trait evolution is challenging, including our current study. A caveat in our study is that holding density may have impacted our results as control fish were housed in an area within the experimental tank that predators could not access. Hence, while fish densities were similar in predation and control areas at that start of the experiment, densities decreased in the predation areas only. While fish density can impact growth parameters (Lorenzen and Enberg 2002), we believe that any such effects are likely negligible due to the relatively brief duration (few weeks) during the adult period that fully grown guppies spent in the experimental tanks. We therefore argue that it is most parsimonious that the group differences between predation and control ones we report in this study were driven predominantly by evolutionary effects driven by strong adult predation.

Subsequently, we need to consider the mechanism that may explain trait differences between our control and predation lines. Was it the non-random change of allele frequency in the predation groups (evolution), phenotypic plasticity, transgenerational plasticity (parental effects), or a mix of those? As we studied fish that experienced predation differences themselves, and the offspring of fish experiencing predation differences, we cannot disentangle those effects, but we may speculate on their relative importance. While three generations may not seem a long period on evolutionary scale, guppies seem to evolve remarkably fast - at a rate of up to seven orders of magnitude faster than in a fossil record (Reznick et al. 1997). Indeed, several independent artificial selection experiments in guppies revealed striking differences after only two to four generations of selection on both morphology and behaviour (selection on relative brain size: 9% after two generations (Kotrschal et al. 2013a), relative telencephalon size: 10.1% in females and 9.5% in males after four generations (Fong et al. 2021), schooling behaviour: 15% after three generations (Kotrschal et al. 2020)). Mesocosm and transplant studies show life history changes in guppies after 4-11 years (Reznick et al. 1997; Reznick and Bryga 1987; Reznick, Bryga, and Endler 1990) and differences in colouration between populations after two years (Endler 1980). Thus, differences manifesting after three generations in our study are in line with previous experimental approaches to guppy evolution. Yet, it is impossible to disentangle whether these differences in our study are due to genetic effects or due to plastic and/or parental effects. Previous research shows that both parental effects and phenotypic plasticity in response to predation can greatly alter an animal’s phenotype (Agrawal, Laforsch, and Tollrian 1999; Moczek et al. 2011; Storm and Lima 2010). For instance, when *Daphnia cucullata* were exposed to water with predator kairomones, they developed defensive structures (through phenotypic plasticity). Individuals of the next generation of these daphnids, which were never exposed to predator cues themselves, were born with similar defensive structures (through maternal effects). This could be explained by themothers ‘anticipating’ that their offspring would be born in a high-predation risk environment and adjusted offspring phenotype accordingly (Agrawal, Laforsch, and Tollrian 1999). In our study, we think the effect of (transgenerational) plasticity is minimal due to the close matching of what predation and control lines experienced during the predation phase (they were both in the same tanks, controls simply protected from consumption). Yet, while evolutionary processes are the likely driver behind the differences found, future work on fish from these lines bred under common garden conditions will be paramount to disentangle the (transgenerational) plasticity and evolutionary mechanisms.

In conclusion, we identified a range of morphological and life history traits that respond to artificial selection on adult predation survival. Our study allows for causal conclusions on the impact of predation on the evolution of those traits and improves our understanding of how life history and morphology evolves. Furthermore, our findings point to the necessity for future research to investigate the complex interactions among various selective agents in shaping these evolutionary processes.

## Materials and methods

### Experimental design

We examined the selective effect of direct adult predation on reproductive and morphological traits by comparing experimental lines of fish (*Poecilia reticulata*) subjected to divergent predation pressures over three generations.

### Selection for predator survival

We used laboratory descendants of Trinidadian guppies collected from high-predation populations from the Quare River in 2005. Stock populations were kept in several large tanks for 14 years, without predators, at Trondheim University. In 2010, 150 animals were brought from Trondheim to Stockholm to start an initial stock population (Vega-Trejo et al., 2022). In 2018, more animals were brought from Trondheim to Stockholm to set up 110 breeding pairs (F0), from which juveniles for three replicates were produced (F1). These juveniles were supplemented with 100 juveniles from the initial stock population (F1). While our population originated from a high-predation locality, we assume that after 14 years of relaxed selection with a species known for its fast evolution (Reznick et al. 1997), we were starting with a low-predation population.

The chosen juveniles were kept in 4L tanks until their sex could be identified (females by their gravid spot, and males by the presence of a modified anal fin called a gonopodium). Mature individuals were kept in groups of 50 individuals in single-sex 50 L tanks until the start of the experiment. For each replicate, a total of 212 mature individuals of each sex were used; of those, 180 were randomly allocated to the predation treatment, 32 to the control treatment. Those fish were transferred to experimental tanks for the selection procedure. Each treatment, replicate, and sex combination was placed in an individual tank. Tank dimensions were 120×110×70 cm, filled to 20 cm with water, for the first two selection events, performed at Stockholm University. For the third selection event performed at Wageningen University, the experimental tank was cylindrical (d = 118 cm, h = 100 cm; filled to 20 cm with water). For all selection events, the bottom was layered with multicoloured limestone gravel (3-8 mm grain size) with which we crafted areas of different depths, so that the water depth ranged from 5-17 cm (15-3 cm gravel) (Figure 5). The shallow area provided refuge for the guppies where the predator could not hunt. One major guppy predator, a pike cichlid (*Chrenicicla alta*) was placed in each tank at the deepest area of the tank and was provided a shelter. The cichlids used were acquired through the aquarium trade and fed with live guppies prior to the experiment. The predation fish (180 individuals) were added to the experimental tanks and were free to roam the entirety of the experimental tank. Control fish (32 individuals) were held in two 11L transparent tanks submerged in the experimental tank (16 individuals per tank). We installed two Eheim filter pumps (60L × h^-1^ per pump) outside the 11L tanks and directed the water flow into each of these tanks to provide olfactory cues for the control fish. Thus, the control fish had visual exposure to the predator and to the behaviour and density of the guppies in the full tank. Additionally, they were exposed to the same water as the predation treatment.

**Figure 5.**
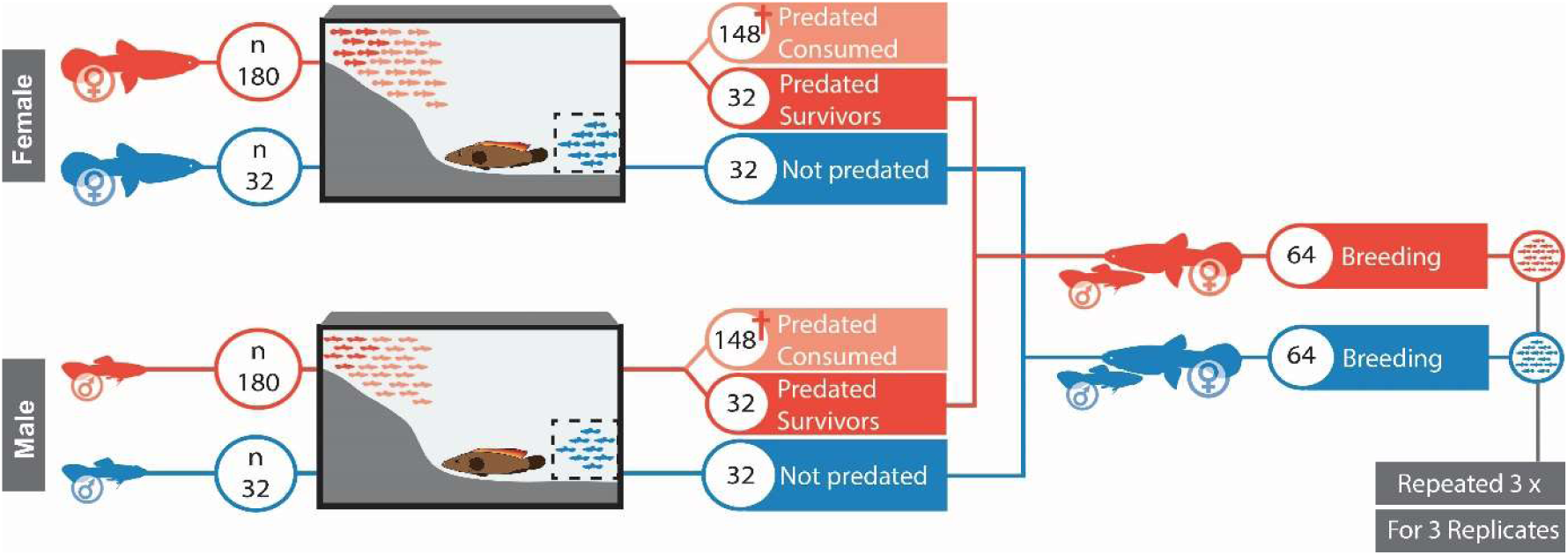
Experimental procedure for the selection for predator survival. Fish from the predation treatment (red) were allowed to swim freely in the tank (N = 180), whereas control fish (blue) were placed in two 11L transparent tanks (N = 16 each, shown in dashed squared) to provide visual and olfactory cues. A predatory cichlid was placed at the deepest area. The sexes and replicates were kept in separate tanks. Of the 180 predation fish introduced, we extracted 32 survivors, as well as 32 control fish. A male and female from the same replicate were matched in breeding pairs to create the next generation. This cycle was repeated three times.

During the first two days of the experiment, we placed a mesh enclosure around the cichlid so predation guppies could escape easily, acclimatize, and learn about the position and potential danger of the predator. This mesh was removed on the third day. We visually monitored the number of fish in the tanks daily. When the number of surviving fish seemed to have reached our target (15% of the group, ∼ 30 individuals), they were captured with a net to be counted. If more than the desired number of survivors was counted, they were returned to the tank. Due to logistical constraints, the final percentage of survivors varied from 13 to 23% between replicates and over the generations. The number of days fish were in the treatment varied from 10 to 52 days for males, and from 12 to 100 days for females. See Supplementary material for details. Because one predator showed signs of stress (hiding and very little feeding) during the first selection experiment, it was replaced with another cichlid after 28 days. Fish were kept at 24 °C under a 12:12 light:dark cycle and fed flake food daily and freshly hatched brine shrimp at least three times per week. Once the selection experiment was completed, a male and a female from the same replicate and treatment were paired up to produce fry. Individuals were placed in a 4L tank with a mesh in the front to allow fry shelter. Tanks were then checked daily (or every other day due to logistical constraints) and the number of fry and date of birth was noted. Fish from the first two generations (F1 and F2) were allowed to breed between 15 and 24 weeks. Note that the variation on time allowed to breed depended on the number of fry produced, as a minimum number of offspring were needed for the subsequent selection procedure. For the last generation (F3), all breeding pairs were allowed to breed for 37 weeks to maximize data collection of their reproductive traits. Once the target number of fry was reached, breeding pairs were euthanised with an overdose of benzocaine and fixed with 4% formalin in buffered phosphate buffer saline (PBS) solution. Once the progeny from each line were fully mature, the selection method was performed again on the next two generations, providing a total of three generations of selection (Figure 5).

### Changes in reproductive traits

Once the survival experiment for both sexes from the same treatment and replicate were completed, a male and female were paired up to produce fry. First, we measured the standard length of both female and male fish by taking individual photos from above with a Nikon DSLR camera. Fish were lit from underneath and photographed from above to enhance contrast. From each photo, we measured standard length with ImageJ(Schneider, Rasband, and Eliceiri 2012). Next, breeding pairs were placed in a 4L tank with mesh in the front to allow fry shelter. Once the breeding pairs were established, we checked tanks daily (or every other day due to logistical constraints) and we recorded the date of birth, treatment, replicate, breeding pair number, and number of fry. We allowed the fish to breed for 37 weeks. From this data, the mean number of fry per breeding pair, total number of fry, number of broods, time to first brood, and interbrood interval could be extrapolated. After breeding and data collection, parental fish were euthanised with an overdose of 2-phenoxyethanol and fixed with 4% formalin in buffered phosphate buffer saline (PBS) solution.

New-born juveniles (F4) were collected on Mondays and Tuesdays for 3 consecutive weeks. Clutches of three or more fry were selected from all different breeding pairs, six groups for each replicate and treatment. Of these selected clutches, three juveniles were randomly selected and photographed on the day of discovery with a Nikon DSLR camera equipped with a Tamron 90mm macro lens. Fry were lit from underneath and photographed from above to enhance contrast. From each photo, we measured standard length with ImageJ.

### Changes in morphological traits

We took photos of approximately 150 adults (F4) per sex and treatment spread out over the three replicates (female predation: 150, female control: 151; male predation: 151, male control: 152). Fifty fish were randomly selected from a single-sex stock tank (45×42×30 cm). These fish were anaesthetized in 4L tanks with water and 2-phenoxyethanol (0.25 mL/L).

Once the fish lost equilibrium, the fish were weighed dry. The fish was then transferred to a petri dish and a photo was taken with a Nikon DSLR camera attached to a tripod. Next, the fish was allowed to recover in fresh water with a large oxygen supply. From the photo, we measured the fish’s standard length, body area, tail length, and eye size using a graphic pad (Wacom Intuos M) and ImageJ. Additionally, for the males, we measured gonopodium length, and colouration (area of black, area of orange, and area of blue iridescence).

Experiments were conducted at Stockholm University, Sweden (November 2019 – December 2020) and Wageningen University, The Netherlands (January 2021 – January 2023).

### Statistical analyses

#### Changes in reproductive traits

We first tested whether the likelihood of producing offspring varied between predation fish and control fish from using a generalised linear model (GLM, binomial distribution) using lme4 (version 1.1-35.1; Bates et al., 2005), with Treatment (Control vs Predation) and Replicate (three levels) as fixed factors.

We then evaluated whether the number of offspring differed between treatments using a GLM (Poisson distribution) including all broods. The model included the interaction between Treatment:Brood, Treatment, Brood, Replicate as fixed factors, and ID as a random factor. We also added Standard length of the mother as a covariate.

To test whether the total amount of offspring produced differed between the treatments, we ran a GLM (Poisson distribution) with Treatment and Replicate as fixed factors, and the Standard length of the mother as a covariate. Similarly, we evaluated with the same factors whether the amount of broods produced over a set amount of time differed between the two treatments with a GLM (Poisson distribution).

To test whether the interbrood interval differed between the treatments we ran two linear models: one for the time to produce the first brood (GLM - Poisson distribution, including Treatment, and Replicate as fixed factors), and another one for interbrood interval for fish that had more than one brood (GLM - Poisson distribution, including Treatment:Brood, Treatment, and Replicate as fixed factors; and ID as a random factor). For both models, the Standard length of the mother was added as a covariate.

#### Changes in morphological traits

To test for differences in standard length of new-born juveniles, we ran a linear model with Treatment and Replicate as fixed factors, the breeding pair from which they were born as a random factor, and the Standard length of the mother as a covariate.

For adults – both male and female – we tested for differences in standard length with fixed factors Treatment, and Replicate. To test for differences in all additional adult female and male traits (body area, weight, tail length, eye size), we ran linear models with fixed factors of Treatment and Replicate, and Standard length as a covariate. For the adult males, we additionally tested for differences in gonopodium size, and black, orange, and iridescence colouration with the same factors as mentioned above.

All statistical analyses were performed in R v.4.3.2 (R Core Team 2023). The model results and code are available in the Supplementary Material.

## Ethics

This research was approved by the Stockholm Ethical Board (Dnr: 11627-2019) and the Netherlands central commission for animal experiments (AVD 10400202010625).

## Acknowledgements

We thank the staff from Carus and the BHE group at WUR for technical and husbandry assistance.

## Author contributions

Conceptualization: AK, RVT

Methodology: HDW, RVT, CVP, AK, JN

Investigation: HDW, RVT, CVP, AK, JN, EP

Writing – original draft: HDW

Writing – review & editing: HDW, RVT, CVP, BP, AK, JN, EP

## Data and materials availability

All data needed to evaluate the conclusions in the paper are present in the paper and/or the Supplementary Materials.

