## Supplementary tables for "Artificial selection for adult predation survival impacts life history and morphology in guppies (*Poecilia reticulata*)"

### Supplemental information

**Table S1.** Outcomes of statistical models for the a) likeliness to breed, b) offspring number, c) total offspring number, d) amount of broods, e) time to produce first brood f) interbrood interval; Nind = number of individuals; Nobs = number of observations; Significant values are given in bold.

| Reproductive traits | Estimate | Standard Error | Z-value | P-value |
|---|---|---|---|---|
| a) Likeliness to breed - Nind = 191 (P: 96, C: 95); Nobs = 191 (P: 96, C: 95) | | | | |
| Intercept | 3.434 | 1.016 | 3.380 | **< 0.001** |
| Treatment | 18.610 | 4544 | 0.004 | 0.997 |
| Replicate2 | 2.157 e-15 | 1.437 | 0.000 | 1.000 |
| Replicate3 | 18.230 | 5517 | 0.003 | 0.997 |
| b) Offspring number - Nind = 191 (P: 96, C: 95); Nobs = 863 (P: 430, C: 433) | | | | |
| Intercept | 1.403 | 0.355 | 3.957 | **< 0.001** |
| Treatment | 0.191 | 0.077 | 2.461 | **0.014** |
| Brood | -0.064 | 0.015 | -4.335 | **< 0.001** |
| Replicate2 | 0.074 | 0.044 | 1.668 | 0.095 |
| Replicate3 | 0.165 | 0.048 | 3.468 | **< 0.001** |
| Stand length mother | 0.024 | 0.013 | 1.835 | 0.067 |
| Treatment:brood | -0.069 | 0.021 | -3.247 | **< 0.001** |
| c) Total offspring number - Nind = 191 (P: 96, C: 95); Nobs = 188 (P: 95, C: 93) | | | | |
| Intercept | 2.888 | 0.499 | 5.792 | **< 0.001** |
| Treatment | -0.006 | 0.073 | -0.076 | 0.940 |
| Replicate2 | 0.114 | 0.062 | 1.837 | 0.066 |
| Replicate3 | 0.231 | 0.067 | 3.425 | **< 0.001** |
| Stand length mother | 0.018 | 0.019 | 0.965 | 0.334 |
| d) Amount of broods - Nind = 191 (P: 96, C: 95); Nobs = 188 (P: 95, C: 93) | | | | |
| Intercept | 1.768 | 0.693 | 2.549 | **0.011** |
| Treatment | 0.002 | 0.102 | 0.020 | 0.984 |
| Replicate2 | 0.050 | 0.088 | 0.567 | 0.571 |
| Replicate3 | 0.079 | 0.094 | 0.843 | 0.399 |
| Stand length mother | -0.011 | 0.026 | -0.409 | 0.683 |
| e) Time to produce first brood - Nind = 191 (P: 96, C: 95); Nobs = 188 (P: 95, C: 93) | | | | |
| Intercept | 4.061 | 0.202 | 20.155 | **< 0.001** |
| Treatment | -0.021 | 0.029 | -0.710 | 0.478 |
| Replicate2 | -0.023 | 0.025 | -0.928 | 0.353 |
| Replicate3 | -0.013 | 0.027 | -0.471 | 0.638 |
| Stand length mother | -0.002 | 0.008 | -0.206 | 0.836 |
| f) Interbrood interval - Nind = 191 (P: 96, C: 95); Nobs = 863 (P: 430, C: 433) | | | | |
| Intercept | 3.864 | 0.196 | 19.669 | **< 0.001** |
| Treatment | -0.050 | 0.046 | -1.092 | 0.275 |
| Replicate2 | -0.071 | 0.025 | -2.894 | **0.004** |
| Replicate3 | -0.056 | 0.026 | -2.133 | **0.033** |
| brood | -0.083 | 0.009 | -9.280 | **< 0.001** |
| Stand length mother | 0.006 | 0.007 | 0.757 | 0.449 |
| Treatment:brood | 0.012 | 0.013 | 0.955 | 0.340 |

**Table S2.** Outcomes of statistical models for the a) standard length, b) weight, c) body area, d) tail length, e) eye size, f) gonopodium length, g) black, h) orange, i) iridescence; Nind = Nobs = number of individuals; Significant values are given in bold.

| Morphological traits | Estimate | Standard Error | Z-value | P-value |
|---|---|---|---|---|
| Juveniles | | | | |
| a) Standard length - Nind = 109 (P: 54, C: 54) | | | | |
| Intercept | 8.708 | 0.898 | 9.693 | **< 0.001** |
| Treatment | 0.396 | 0.141 | 2.804 | **0.005** |
| Replicate2 | -0.248 | 0.115 | -2.152 | **0.031** |
| Replicate3 | -0.248 | 0.104 | 0.474 | 0.635 |
| Stand length mother | -0.038 | 0.035 | -1.107 | 0.268 |
| Females | | | | |
| a) Standard length - Nind = 301 (P: 150, C: 151) | | | | |
| Intercept | 27.894 | 0.249 | 112.030 | **< 0.001** |
| Treatment | -1.432 | 0.250 | -5.740 | **< 0.001** |
| Replicate2 | 0.291 | 0.305 | 0.950 | 0.341 |
| Replicate3 | -0.332 | 0.307 | -1.080 | 0.279 |
| b) Weight - Nind = 301 (P: 150, C: 151) | | | | |
| Intercept | -374.278 | 79.970 | -4.680 | **< 0.001** |
| Treatment | -395.890 | 113.197 | -3.497 | **< 0.001** |
| Replicate2 | 32.235 | 10.976 | 2.937 | **< 0.001** |
| Replicate3 | 58.343 | 11.012 | 5.298 | **0.003** |
| Stand length | 28.420 | 2.874 | 9.890 | **< 0.001** |
| Treatment:stand length | 15.562 | 4.156 | 3.744 | **< 0.001** |
| c) Body area - Nind = 301 (P: 150, C: 151) | | | | |
| Intercept | -80.503 | 13.019 | -6.184 | **< 0.001** |
| Treatment | 5.809 | 2.089 | 2.781 | **0.005** |
| Replicate2 | 3.921 | 2.426 | 1.616 | 0.106 |
| Replicate3 | -23.540 | 2.440 | -9.649 | **< 0.001** |
| Stand length | 8.381 | 0.461 | 18.168 | **< 0.001** |
| d) Tail length - Nind = 301 (P: 150, C: 151) | | | | |
| Intercept | 2.485 | 0.383 | 6.488 | **< 0.001** |
| Treatment | 0.047 | 0.062 | 0.771 | 0.441 |
| Replicate2 | 0.023 | 0.072 | 0.323 | 0.747 |
| Replicate3 | 0.287 | 0.072 | 3.973 | **< 0.001** |
| Stand length | 0.209 | 0.013 | 15.405 | **< 0.001** |
| e) Eye size - Nind = 301 (P: 150, C: 151) | | | | |
| Intercept | 0.626 | 0.252 | 2.484 | **0.013** |
| Treatment | -0.035 | 0.040 | -0.873 | 0.383 |
| Replicate2 | -0.056 | 0.047 | -1.197 | 0.231 |
| Replicate3 | -0.023 | 0.047 | -0.491 | 0.624 |
| Stand length | 0.112 | 0.009 | 12.576 | **< 0.001** |
| Males | | | | |
| a) Standard length - Nind = 303 (P: 151, C: 152) | | | | |
| Intercept | 20.922 | 0.142 | 146.830 | **< 0.001** |
| Treatment | -0.045 | 0.142 | -0.320 | 0.752 |
| Replicate2 | -0.701 | 0.174 | -4.020 | **< 0.001** |
| Replicate3 | -0.218 | 0.174 | 1.251 | 0.211 |
| b) Weight - Nind = 303 (P: 151, C: 152) | | | | |
| Intercept | -280.334 | 23.549 | -11.904 | **< 0.001** |
| Treatment | 45.019 | 31.476 | 1.430 | 0.153 |
| Replicate2 | 4.544 | 2.419 | 1.879 | 0.060 |
| Replicate3 | 3.408 | 2.341 | 1.456 | 0.146 |
| Stand length | 21.428 | 1.134 | 18.894 | **< 0.001** |

| | | | | |
|---|---|---|---|---|
| Treatment:stand length | -2.485 | 1.526 | -1.629 | 0.103 |
| c) Body area - Nind = 303 (P: 151, C: 152) | | | | |
| Intercept | -75.483 | 4.252 | -17.750 | < 0.001 |
| Treatment | -0.511 | 0.499 | -1.020 | 0.306 |
| Replicate2 | -1.243 | 0.629 | -1.980 | 0.0479 |
| Replicate3 | -1.355 | 0.612 | -2.210 | 0.0269 |
| Stand length | 7.618 | 0.202 | 37.740 | < 0.001 |
| d) Tail length - Nind = 303 (P: 151, C: 152) | | | | |
| Intercept | 1.952 | 0.854 | 2.285 | 0.022 |
| Treatment | -0.355 | 0.099 | -3.577 | < 0.001 |
| Replicate2 | -0.054 | 0.125 | -0.436 | 0.663 |
| Replicate3 | -0.120 | 0.122 | -0.978 | 0.328 |
| Stand length | 0.264 | 0.041 | 6.504 | < 0.001 |
| e) Eye size - Nind = 303 (P: 151, C: 152) | | | | |
| Intercept | -0.509 | 0.219 | -2.323 | 0.020 |
| Treatment | 0.047 | 0.026 | 1.843 | 0.065 |
| Replicate2 | -0.017 | 0.032 | -0.513 | 0.608 |
| Replicate3 | -0.067 | 0.032 | -2.115 | 0.034 |
| Stand length | 0.137 | 0.010 | 13.167 | < 0.001 |
| f) Gonopodium length - Nind = 303 (P: 151, C: 152) | | | | |
| Intercept | 3.497 | 0.431 | 8.122 | < 0.001 |
| Treatment | -0.487 | 0.050 | -9.717 | < 0.001 |
| Replicate2 | -0.045 | 0.064 | -0.702 | 0.483 |
| Replicate3 | -0.031 | 0.062 | -0.506 | 0.613 |
| Stand length | 0.037 | 0.020 | 1.826 | 0.068 |
| g) Black - Nind = 303 (P: 151, C: 152) | | | | |
| Intercept | -15.145 | 3.858 | -3.926 | < 0.001 |
| Treatment | -0.148 | 0.453 | -0.327 | 0.743 |
| Replicate2 | 1.203 | 0.570 | 2.109 | 0.035 |
| Replicate3 | 1.000 | 0.556 | 1.799 | 0.072 |
| Stand length | 1.346 | 0.183 | 7.350 | < 0.001 |
| h) Orange - Nind = 303 (P: 151, C: 152) | | | | |
| Intercept | -1.761 | 2.907 | -0.606 | 0.545 |
| Treatment | 0.322 | 0.341 | 0.943 | 0.346 |
| Replicate2 | 1.307 | 0.430 | 3.043 | 0.002 |
| Replicate3 | 0.350 | 0.419 | 0.837 | 0.403 |
| Stand length | 0.273 | 0.138 | 1.980 | 0.048 |
| i) Iridescence - Nind = 303 (P: 151, C: 152) | | | | |
| Intercept | -7.097 | 4.178 | -1.699 | 0.089 |
| Treatment | -0.128 | 0.490 | -0.260 | 0.795 |
| Replicate2 | -2.114 | 0.618 | -3.423 | < 0.001 |
| Replicate3 | -1.424 | 0.602 | -2.367 | 0.018 |
| Stand length | 0.654 | 0.198 | 3.298 | < 0.001 |

**Table S3.** Number of days in the experimental treatment for each generation and replicate.

| | | Replicate 1 | Replicate 2 | Replicate 3 |
|---|---|---|---|---|
| F1 | | | | |
| | Males | 52 | 40 | 22 |
| | Females | 77 | 90 | 100 |
| F2 | | | | |
| | Males | 19 | 10 | 21 |
| | Females | 19 | 12 | 14 |
| F3 | | | | |
| | Males | 45 | 49 | 36 |
| | Females | 85 | 59 | 71 |
